# BglC of *Streptomyces scabiei* is a scopolin-hydrolyzing β-glucosidase interfering with the host defense mechanism

**DOI:** 10.1101/2021.10.20.465124

**Authors:** Benoit Deflandre, Sébastien Rigali

## Abstract

The beta-glucosidase BglC fulfills multiple functions in both primary metabolism and induction of pathogenicity of *Streptomyces scabiei*, the causative agent of the common scab disease of root and tuber crops. Indeed, this enzyme hydrolyzes cellobiose and cellotriose to directly feed glycolysis with glucose, but also modifies the intracellular concentration of these cello-oligosaccharides which are the virulence elicitors. The inactivation of *bglC* also led to unexpected phenotypes such as the constitutive overproduction of thaxtomin A, the main virulence determinant of *S. scabiei*. In this work we revealed a new target substrate of BglC, the phytoalexin scopolin. Removal of the glucose moiety of scopolin generates scopoletin, a potent inhibitor of thaxtomin A production. The hydrolysis of scopolin by BglC presents substrate inhibition kinetics which contrasts with the typical Michaelis–Menten saturation curve previously observed for the degradation of its natural substrate cellobiose. Our work therefore reveals that BglC targets both cello-oligosaccharide elicitors emanating from the hosts of *S. scabiei*, and the scopolin phytoalexin generated by the host defense mechanisms, thereby occupying a key position to fine-tune the production of the main virulence determinant thaxtomin A.

## Introduction

The common scab disease on root and tuber crops is caused by a dozen of bacterial strains belonging to the Gram-positive *Streptomyces* genus, with strain *Streptomyces scabiei* (syn. *S. scabies*) as model species. The major virulence determinants are 4□nitroindol□3□yl□containing 2,5□dioxopiperazines called thaxtomins, which are potent cellulose synthesis inhibitors in higher plants [1]. Thaxtomin A is the most abundant toxin and its biosynthesis, together with many other specialized metabolites that compose the “virulome” of *S. scabiei* [2–5], is triggered by cello-oligosaccharides with cellotriose being the main elicitor emanating from the plant hosts [6–8]. As virulence elicitor, cellotriose is imported by the ABC-type transporter CebEFG-MsiK [6], and, once inside the cytoplasm, it prevents - together with cellobiose generated from cellobiose (see below) - DNA-binding of the transcriptional repressor CebR which allows the expression of the thaxtomin regulatory and biosynthetic genes [9].

One gene/protein intimately linked to the other players involved in the perception and transport of cello-oligosaccharides suggests that our current understanding of the virulence signaling pathway is partial. The piece that renders the jigsaw more complicated than initially thought is the beta-glucosidase BglC, whose primary function is to feed glycolysis with glucose released from the hydrolysis of cellotriose and cellobiose [10] (Figure 1, role ⍰). However, complete degradation of cellotriose and cellobiose into glucose by BglC may prevent *S. scabiei* to accumulate the elicitors of the virulome and behave like a saprophyte. How does BglC manage to control the intracellular concentration of cello-oligosaccharides for determining the proper timing of the pivotal switch from the saprophytic to the pathogenic lifestyle is currently unknown (Figure 1, role ⍰). Moreover, the deletion of *bglC* led to an unpredicted phenotype where *S. scabiei* (strain Δ*bglC*) constitutively produces the thaxtomin phytotoxins, in culture conditions not supplied with cello-oligosaccharides, thereby displaying an hypervirulent phenotype [10]. This unexpected cello-oligosaccharide-independent overproduction of thaxtomin A in strain Δ*bglC* suggests that, in the wild-type strain *S. scabiei* 87-22, BglC must repress a yet unknown alternative path to pathogenicity (Figure 1, role ⍰). Finally, we discovered that the primary function of *bglC*/BglC, *i*.*e*., the utilization of cello-oligosaccharides, is safeguarded by a mechanism of genetic compensation that awakens the expression of alternative beta-glucosidases upon perceiving the loss of *bglC* [11]. When BglC is active, the transcription of these alternative beta-glucosidases is kept to a minimal level and is not induced by cello-oligosaccharides (Figure 1, role ^17^).

**Figure 1.**
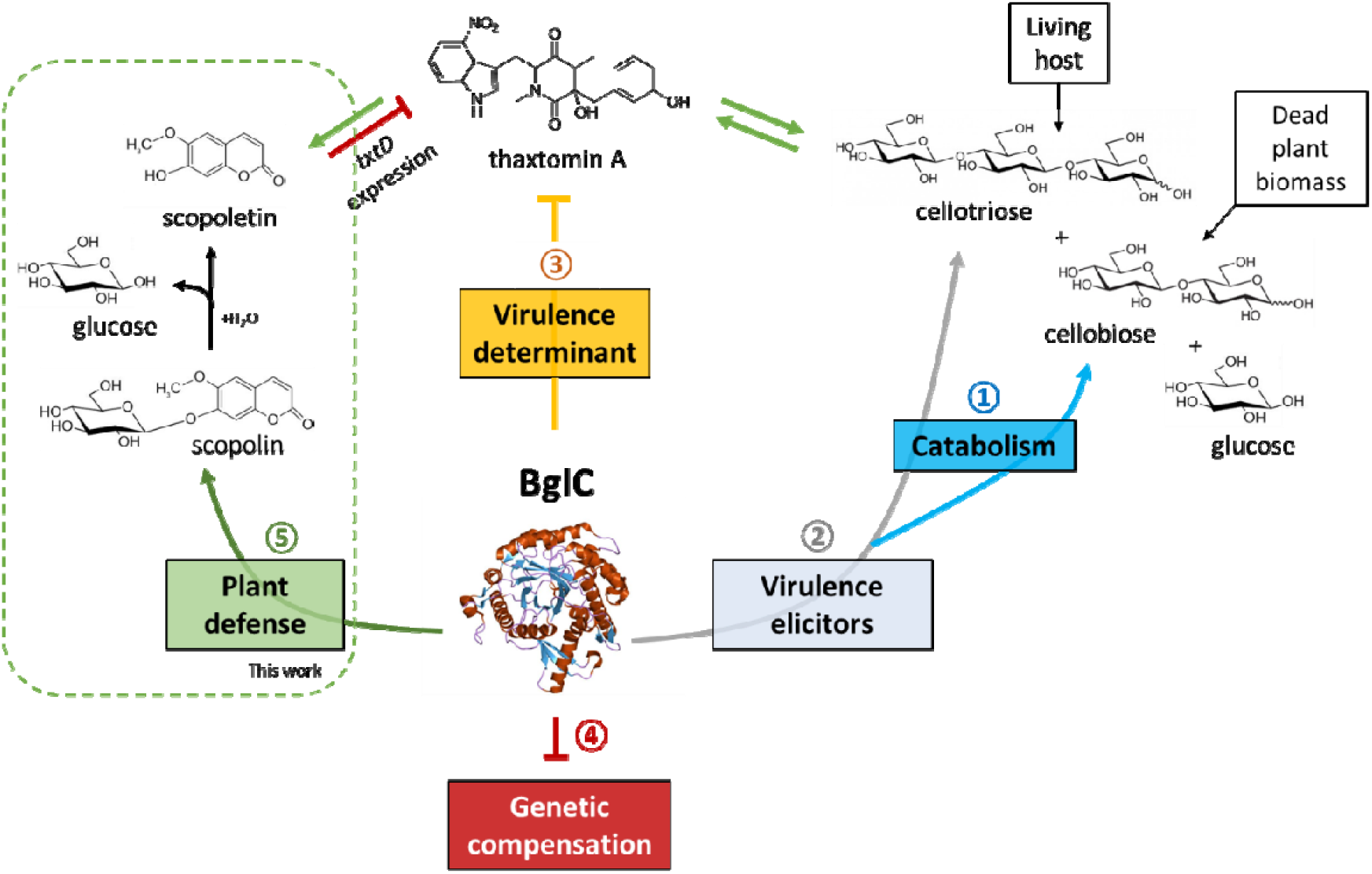
The five biological functions associated with BglC of *S. scabiei* 87-22. ⍰The primary function of BglC is to hydrolyze cellobiose - the main byproduct resulting from the degradation of cellulose from decaying plant material – and cellotriose to directly feed glycolysis with glucose [10]; ⍰BglC degrades cellotriose as virulence elicitor released from living hosts [7] and therefore fine-tunes the strength of the onset of the pathogenic lifestyle of *S. scabiei*; ⍰BglC represses the production of thaxtomin A, the main virulence determinant of *S. scabiei*, by a yet unknown cello-oligosaccharide-independent mechanism [10]; ⍰The beta-glucosidase activity of BglC is safeguarded by a mechanism of genetic compensation that represses the expression/production of alternative beta-glucosidases [11]; ⍰In this work, we showed that BglC also hydrolyzes the phytoalexin scopolin into scopoletin, a strong inhibitor of thaxtomin A produced by plants colonized by *S. scabiei* [12].

For its multiple roles as well as the unexpected phenotypes and physiological responses of the *bglC* null mutant, BglC is an untapped source for research themes likely to provide crucial information for properly understanding the lifestyle of *S. scabiei*. In this work, we unveil a fifth role for BglC, *i*.*e*., associated with a defense mechanism of its hosts as this beta-glucosidase can also hydrolyze the phytoalexin scopolin into scopoletin, the latter being a strong thaxtomin A inhibitor which is produced by plants upon colonization by *S. scabiei*.

## Materials and methods

### Heterologous production of His_6_-tagged proteins and purification

BglC of *S. scabiei* 87-22 with a six-histidine tag fused to the N-terminus part of the protein (His_6_-BglC) was produced in *Escherichia coli* BL21(DE3) Rosetta™ and purified by nickel affinity chromatography as already described in [10,11]. The pure protein was stored and used in HEPES buffer (50 mM, pH 7.5).

### TLC for hydrolysis of carbohydrates

Semi-quantitative substrate degradation was assessed by thin layer chromatography (TLC). Reactions were carried out with His_6_-BglC (1 μM) and the substrates (5 mM) in HEPES 50 mM pH 7.5 at 40°C for 10 minutes. At the end of the reaction, the mixture was incubated for 5 minutes in a boiling water bath to inactivate the enzyme. 1-μL samples of the inactivated reaction mixtures were spotted next to undigested standards on aluminum-backed TLC plates (Silica gel Matrix, Sigma-Aldrich) and thoroughly dried. The protocol, adapted from [13], consisted in eluting the loaded TLC plate in a TLC chamber filled with an elution buffer (Chloroform – Methanol – Acetic acid – Water (50:50:15:5 (v/v))). After air-drying the eluted plate, sulfuric acid (5%) in ethanol was sprayed onto the TLC plate, and the excess liquid was drained. The revelation was finally conducted by heating the TLC plate.

### Determination of kinetic parameters for His_6_-BglC

The hydrolysis of scopolin was followed by glucose quantification using the D-Glucose HK Assay kit (Megazyme) following the microplate procedure. His_6_-BglC was mixed with scopolin at variable concentrations in HEPES 50 mM pH 7.5, and the incubation was conducted at 40°C for 4 minutes. The reaction was terminated by a 5-minutes incubation in a boiling water bath. At least 10 concentrations – if possible distributed around the K_m_ value – were tested in triplicate to estimate initial velocity values. The obtained data – initial velocity (V_i_, mM/min) in function of scopolin concentration ([S], mM) – were fitted to the Substrate inhibition equation V_i_ =(V_max_*[S])/(K_m_□+□[S]*(1+[S]/K_i_)) using the GraphPad Prism (version 9.2.0) software.

## Results

Due to the multiple functions fulfilled by BglC that cannot be explained by a single activity on its natural substrates cellobiose and cellotriose, we investigated if this beta-glucosidase would target other carbohydrates with a terminal glucose attached by a β-1,4 linkage. BglC of *S. scabiei* 87-22 was suggested by KEGG pathway [14] as candidate beta-glucosidase possibly involved in cyanoamino acid metabolism (https://www.genome.jp/kegg-bin/show_pathway?scb00460+SCAB_57721). Two cyanogenic glucosides were tested as possible targets of BglC, *i*.*e*., amygdalin and linamarin. The glycone moiety of amygdalin is the disaccharide gentiobiose (also called amygdalose) and its aglycone part is mandelonitrile, the cyanohydrin of benzaldehyde; linamarin is a glucoside of acetone cyanohydrin. According to the hypervirulent phenotype of the *bglC* null mutant [10], the scopolin heteroside was additionally selected as possible substrate of BglC. Scopolin is the glucoconjugate of scopoletin, a phytoalexin produced by plants under colonization by *S. scabiei* or under application of pure thaxtomin A. The production of scopoletin and its glucoconjugate (e.g. scopolin) was observed in *Arabidopsis thaliana* seedlings, tobacco leaves [12], and in potato tuber tissue under pathogen invasion [15,16]. Lerat and colleagues observed that the production of scopoletin under *S. scabiei* infection appears to be responsive to thaxtomin A and this compound acts as a powerful inhibitor of the expression of *txtD*, consequently limiting thaxtomin biosynthesis [12].

Semi-quantitative substrate degradation was first assessed by thin layer chromatography (TLC) where amygdalin and its glucoside moiety gentiobiose, linamarin, and scopolin (5 mM) were incubated with 1 μM pure six-histidine tagged BglC (His_6_-BglC) (see the Materials and methods section for detailed protocols). Cellobiose was also included in these degradation assays as positive control to assess the correct activity of pure His_6_-BglC. Reaction samples were spotted on TLC plates and migrated in an elution chamber to separate the glucose moiety (or possibly the gentiobiose moiety in the case of amygdalin) from the rest of the substrate. As shown in Figure 2, glucose is released from cellobiose when the disaccharide is incubated with His_6_-BglC, thereby proving that the pure beta-glucosidase is active. His_6_-BglC did not degrade neither D-amygdalin nor gentiobiose alone, the glycone moiety of amygdalin. Similarly, glucose was not released from linamarin upon incubation with His_6_-BglC. In contrast, a clear spot with the same migration rate as glucose was observed when scopolin was incubated with His_6_-BglC.

**Figure 2.**
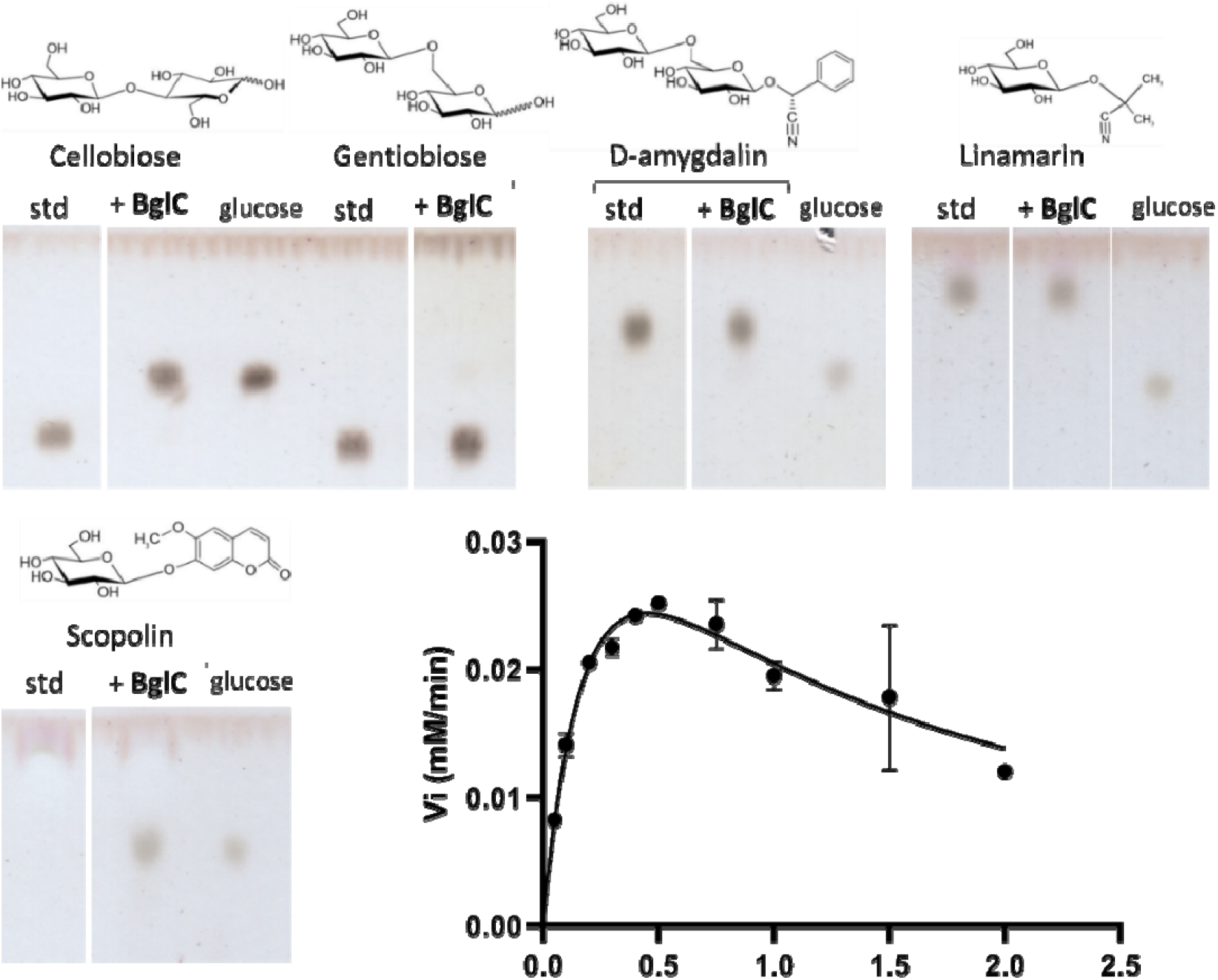
Beta-glucosidase activity of BglC on scopolin. Upper and bottom left panels. TLC plates revealing the release of glucose after incubation of a variety of substrates (5 mM) with His_6_-BglC (1 μM) compared to the intact substrate (standard (std)). The respective chemical structures of the substrates are displayed above each name. Bottom right panel. Plot of the initial velocity (V_i_, mM/min) estimated by the rate of glucose released by His_6_-BglC as a function of scopolin concentrations (in mM). Individual values were entered into the GraphPad Prism software (9.2.0) which fitted the data to the Substrate Inhibition model. Error bars display the standard deviation values determined for the V_i_ for three replicates at each substrate concentration.

Following these preliminary enzymatic assays on TLC, the hydrolysis of scopolin by His_6_-BglC was further investigated by determining the kinetic parameters of the reaction. As deduced by the non-linear regression profile (Figure 2, bottom right panel), His_6_-BglC is subjected to substrate inhibition when scopolin is provided at around 1 mM and higher concentrations. The inhibitory constant (K_i_) for His_6_-BglC was calculated at 0.68 mM as deduced using the equation V_i_ = (V_max_*[S])/(K_m_□+□[S]*(1+[S]/K_i_)) proposed by the GraphPad Prism software (version 9.2.0). The K_m_ (affinity of the enzyme for the substrate) and k_cat_ (turnover of substrate molecules per second) values were 0.30 mM (0.77 mM for cellobiose), and 9.4 s^-1^ (6.7 s^-1^ for cellobiose), respectively.

## Discussion

In this work we revealed that the phytoalexin scopolin is a new substrate hydrolyzed by the beta-glucosidase BglC, further expanding the crucial biological roles played by this enzyme in *S. scabiei* (Figure 1, role ⍰). By hydrolyzing the molecules (cellotriose and cellobiose) that will activate thaxtomin production, as well as a molecule (scopolin) that will instead generate an inhibitor of thaxtomin biosynthesis, BglC occupies a key position to fine-tune the production of the main virulence determinant of *S. scabiei* (Figure 3).

**Figure 3.**
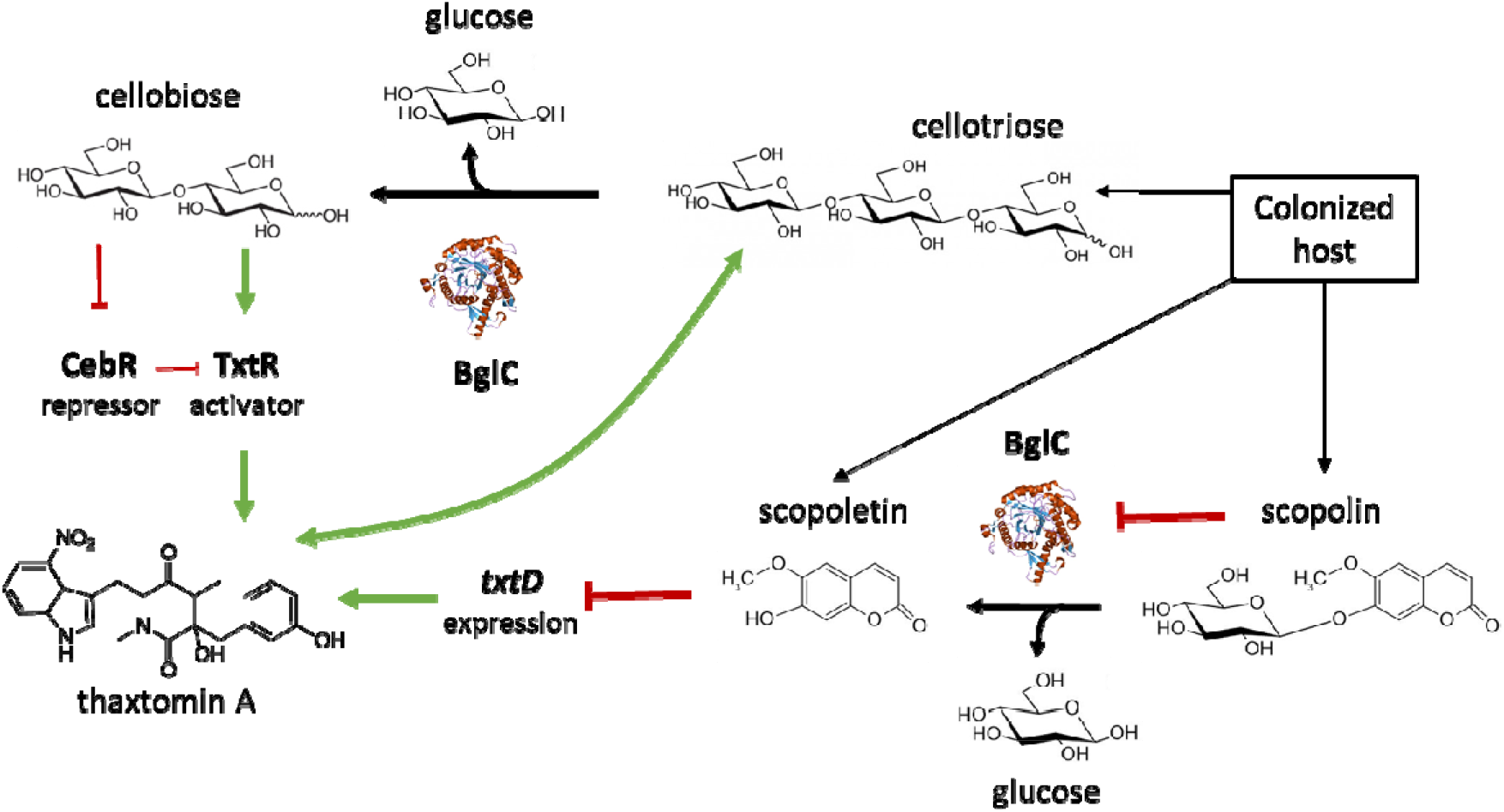
BglC-mediated fine-tuning of thaxtomin production in *S. scabiei*. Cellotriose emanating from expanding plant tissues triggers thaxtomin A production which in turn causes more release of cellotriose from the host [7]. Perception of thaxtomin A by the colonized host induces the production of the scopolin and scopoletin phytoalexins [12], the latter being a strong repressor of thaxtomin A production by reducing the expression of *txtD* (also called *nos, scab31841*) [12]. Hydrolysis of cellotriose by BglC generates cellobiose, the best allosteric effector of CebR - the repressor of the thaxtomin biosynthetic gene cluster - which also directly represses the expression of the thaxtomin pathway-specific activator TxtR [17]. High concentrations of scopolin would inhibit the BglC activity which would result in i) less accumulation of scopoletin (a thaxtomin A inhibitor), and ii) limit the degradation of cellotriose (the thaxtomin A inducer), both consequences presumed to cause increase thaxtomin A production.

The kinetic values for the hydrolysis of cellobiose and scopolin are in the same order of magnitude. The major difference between the two substrates is that for scopolin, the activity of BglC is inhibited at high concentrations of substrate, whereas in the case of cello-oligosaccharides, BglC remains fully active at the highest concentrations and the kinetic analysis follows a classical Michaelis-Menten curve [10]. In the proposed model presented in Figure 3, high concentrations of scopolin would inhibit the activity of BglC which would result in i) less accumulation of scopoletin (a thaxtomin production inhibitor), and ii) limit the degradation of cellotriose (the thaxtomin production inducer), both consequences are presumed to cause increase thaxtomin production.

## Acknowledgments

The work of BD was supported by an aspirant grant from the FNRS (grant 1.A618.18), and a FNRS grant “Crédit de recherche” (grant CDR/OL J.0158.21) to SR. SR is senior research associate of the FRS-FNRS (Brussels, Belgium). We are very grateful to Nudzejma Stulanovic for the careful reading of the manuscript and molecule drawing.

## References

[1] R. Loria, D.R.D. Bignell, S. Moll, J.C. Huguet-Tapia, M.V. Joshi, E.G. Johnson, R.F. Seipke, D.M. Gibson, Thaxtomin biosynthesis: the path to plant pathogenicity in the genus Streptomyces, Antonie Van Leeuwenhoek. 94 (2008) 3–10. https://doi.org/10.1007/s10482-008-9240-4.

[2] J. Liu, L.-F. Nothias, P.C. Dorrestein, K. Tahlan, D.R.D. Bignell, Genomic and Metabolomic Analysis of the Potato Common Scab Pathogen Streptomyces scabiei, ACS Omega. 6 (2021) 11474–11487. https://doi.org/10.1021/acsomega.1c00526.

[3] Y. Li, J. Liu, G. Díaz-Cruz, Z. Cheng, D.R.D. Bignell, Virulence mechanisms of plant-pathogenic Streptomyces species: an updated review, Microbiology (Reading). 165 (2019) 1025–1040. https://doi.org/10.1099/mic.0.000818.

[4] D.R.D. Bignell, J.C. Huguet-Tapia, M.V. Joshi, G.S. Pettis, R. Loria, What does it take to be a plant pathogen: genomic insights from Streptomyces species, Antonie Van Leeuwenhoek. 98 (2010) 179–194. https://doi.org/10.1007/s10482-010-9429-1.

[5] B. Deflandre, N. Stulanovic, S. Planckaert, S. Anderssen, B. Bonometti, L. Karim, W. Coppieters, B. Devreese, S. Rigali, The virulome of Streptomyces scabiei in response to cello-oligosaccharides elicitors, 2021. https://doi.org/10.1101/2021.08.10.455888.

[6] S. Jourdan, I.M. Francis, M.J. Kim, J.J.C. Salazar, S. Planckaert, J.-M. Frère, A. Matagne, F. Kerff, B. Devreese, R. Loria, S. Rigali, The CebE/MsiK Transporter is a Doorway to the Cello-oligosaccharide-mediated Induction of Streptomyces scabies Pathogenicity, Sci Rep. 6 (2016) 27144. https://doi.org/10.1038/srep27144.

[7] E.G. Johnson, M.V. Joshi, D.M. Gibson, R. Loria, Cello-oligosaccharides released from host plants induce pathogenicity in scab-causing Streptomyces species, Physiological and Molecular Plant Pathology. 71 (2007) 18–25. https://doi.org/10.1016/j.pmpp.2007.09.003.

[8] S. Jourdan, I.M. Francis, B. Deflandre, R. Loria, S. Rigali, Tracking the Subtle Mutations Driving Host Sensing by the Plant Pathogen Streptomyces scabies, MSphere. 2 (2017) e00367–16. https://doi.org/10.1128/mSphere.00367-16.

[9] I.M. Francis, S. Jourdan, S. Fanara, R. Loria, S. Rigali, The cellobiose sensor CebR is the gatekeeper of Streptomyces scabies pathogenicity, MBio. 6 (2015) e02018. https://doi.org/10.1128/mBio.02018-14.

[10] S. Jourdan, I.M. Francis, B. Deflandre, E. Tenconi, J. Riley, S. Planckaert, P. Tocquin, L. Martinet, B. Devreese, R. Loria, S. Rigali, Contribution of the β-glucosidase BglC to the onset of the pathogenic lifestyle of Streptomyces scabies, Mol Plant Pathol. 19 (2018) 1480–1490. https://doi.org/10.1111/mpp.12631.

[11] B. Deflandre, N. Thiébaut, S. Planckaert, S. Jourdan, S. Anderssen, M. Hanikenne, B. Devreese, I. Francis, S. Rigali, Deletion of bglC triggers a genetic compensation response by awakening the expression of alternative beta-glucosidase, Biochim Biophys Acta Gene Regul Mech. 1863 (2020) 194615. https://doi.org/10.1016/j.bbagrm.2020.194615.

[12] S. Lerat, A.H. Babana, M. El Oirdi, A. El Hadrami, F. Daayf, N. Beaudoin, K. Bouarab, C. Beaulieu, Streptomyces scabiei and its toxin thaxtomin A induce scopoletin biosynthesis in tobacco and Arabidopsis thaliana, Plant Cell Rep. 28 (2009) 1895–1903. https://doi.org/10.1007/s00299-009-0792-1.

[13] J. Gao, W. Wakarchuk, Characterization of Five β-Glycoside Hydrolases from Cellulomonas fimi ATCC 484, Journal of Bacteriology. 196 (2014) 4103–4110. https://doi.org/10.1128/JB.02194-14.

[14] M. Kanehisa, A database for post-genome analysis, Trends Genet. 13 (1997) 375–376. https://doi.org/10.1016/s0168-9525(97)01223-7.

[15] D.D. Clarke, The accumulation of scopolin in potato tissue in response to infection, Physiological Plant Pathology. 3 (1973) 347–358. https://doi.org/10.1016/0048-4059(73)90006-4.

[16] P. Nolte, G.A. Secor, N.C. Gudmestad, P.J. Henningson, Detection and identification of fluorescent compounds in potato tuber tissue with corky patch syndrome, American Potato Journal. 70 (1993) 649–666. https://doi.org/10.1007/BF02849154.

[17] M.V. Joshi, D.R.D. Bignell, E.G. Johnson, J.P. Sparks, D.M. Gibson, R. Loria, The AraC/XylS regulator TxtR modulates thaxtomin biosynthesis and virulence in Streptomyces scabies, Mol Microbiol. 66 (2007) 633–642. https://doi.org/10.1111/j.1365-2958.2007.05942.x.

